# Machupo virus with mutations in the transmembrane domain and glycosylation sites is attenuated and immunogenic in animal models of Bolivian Hemorrhagic Fever

**DOI:** 10.1101/2021.12.02.471047

**Authors:** Emily K Mantlo, Junki Maruyama, John T Manning, Timothy G Wanninger, Cheng Huang, Jeanon N Smith, Michael Patterson, Slobodan Paessler, Takaaki Koma

**Affiliations:** Department of Pathology, University of Texas Medical Branch at Galveston, Texas, United States of America; Department of Microbiology, Tokushima University Graduate School of Biomedical Sciences, Tokushima, Japan, 770-8503

## Abstract

Several highly pathogenic mammarenaviruses cause severe hemorrhagic and neurologic disease in humans, for which vaccines and antivirals are limited or unavailable. New World (NW) mammarenavirus Machupo virus (MACV) infection causes Bolivian hemorrhagic fever in humans. We previously reported that the disruption of specific N-linked glycan sites on the glycoprotein (GPC) partially attenuate MACV in an IFN-αβ/γ receptor knockout mouse model. However, some capability to induce neurological pathology still remained. Highly pathogenic Junin virus (JUNV) is another NW arenavirus closely related to MACV. A F427I substitution in the GPC transmembrane domain (TMD) rendered JUNV attenuated in a lethal mouse model after intracranial inoculation. In this study, we rationally designed and rescued a MACV containing mutations at two glycosylation sites and the corresponding F438I substitution in GPC TMD. The MACV mutant is fully attenuated in IFN-αβ/γ receptor knockout mice and outbred guinea pigs. Furthermore, inoculation with this mutant MACV fully protected guinea pigs from wild-type MACV lethal challenge. Lastly, we found the GPC TMD F438I substitution greatly impaired MACV growth in neuronal cell lines of mouse and human origins. Our results highlight the critical roles of the glycans and the TMD on the GPC in arenavirus virulence, which informs the rational design of potential vaccine candidates for highly pathogenic arenaviruses.

**Importance:** For arenaviruses, the only vaccine available is the live-attenuated Candid#1 vaccine, a JUNV vaccine approved in Argentina. We and others have found that the glycans on GPC and the F427 residue in the GPC TMD are important for virulence of JUNV. Nevertheless, mutating either of them is not sufficient for full and stable attenuation of JUNV. Using reverse genetics, we disrupted specific glycosylation sites on MACV GPC, and also introduced the corresponding F438I substitution in the GPC TMD. This MACV mutant is fully attenuated in two animal models and protects animals from lethal infection. Thus, our studies highlight the feasibility of rational attenuation of highly pathogenic arenaviruses for vaccine development. Another important finding from this study is that the F438I substitution in GPC TMD could substantially affect MACV replication in neurons. Future studies are warranted to elucidate the underlying mechanism and the implication of this mutation in arenavirus neural tropism.

## Introduction

Highly pathogenic mammarenaviruses can cause severe hemorrhagic fevers in humans. The Old World mammarenavirus, Lassa virus, is responsible for an estimated 300,000 infections and 5,000 deaths per year in West Africa^1^. New World mammarenaviruses, including Junin virus (JUNV), Machupo virus (MACV), Guanarito virus (GTOV), Chapare virus (CHAPV), and Sabia virus (SABV), periodically emerge in parts of South America to cause localized epidemics with high case fatality rates^2, 3^. The two most common New World arenaviruses, JUNV and MACV, are the causative agents of Argentine hemorrhagic fever (AHF) and Bolivian hemorrhagic fever (BHF), respectively. AHF and BHF share a remarkably similar clinical course of illness, with patients progressing from an initial non-specific prodromal phase into a biphasic hemorrhagic and neurologic disease with an overall case fatality rate ranging from 15-35%^4–6^. Hemorrhagic manifestations include epistaxis, petechiae, hypotension, hematemesis, and mucosal hemorrhaging. Neurologic manifestations include tremors, delirium, muscle spasms, and coma^7, 8^. Overall, there are no FDA-approved treatments or vaccines for any of these infections. Thus, there is an urgent need for new countermeasures for highly pathogenic arenaviruses.

The arenavirus genome consists of 4 viral genes encoding 4 viral proteins in a bisegmented, antisense/ambisense RNA genome^9^. The large (L) segment encodes the RNA-dependent RNA polymerase (L) and the small RING-finger matrix protein, Z. The small (S) segment encodes the nucleoprotein (NP) and the glycoprotein complex (GPC). GPC is further cleaved by cellular proteases into stable signal peptide (SSP), GP1, and GP2^10–12^. Arenaviruses enter the cell through receptor-mediated endocytosis, binding to their cellular receptor via GP1, which is the primary target of host neutralizing antibodies^13, 14^. Old World arenaviruses primarily use α-dystroglycan as a receptor^15^, while New World arenaviruses primarily use transferrin receptor 1 (TfR1)^16, 17^. Upon entry into the endosome, a pH-dependent fusion process mediated by GP2 allows for the release of the viral genomic RNAs^18, 19^, where early translation of NP and L protein begins. Later in the replication process, GPC and Z proteins are produced, and GPC is cleaved in the endoplasmic reticulum before traveling to the Golgi, where a series of glycans are post-translationally added to the protein^20–23^. This is followed by protein trafficking to the cell surface, assembly of the viral particles, and release from the cell.

Only one arenavirus vaccine, the live-attenuated JUNV strain, Candid#1 (Cd#1), has been approved only for use in Argentina^24, 25^. This vaccine was created through the serial passaging of a lethal isolate of JUNV in animals and cultured cells, leading to the acquisition of a series of attenuating mutations. The strain was first passaged through guinea pigs twice, followed by 44 passages in suckling mouse brains^25, 26^. During this passaging, the strain acquired a T168A substitution in GP1 that led to the loss of a glycan^22, 23^. After the 44 passages in mouse brain, the resultant strain, XJ44, was attenuated in guinea pigs but was still virulent in suckling mice after intracranial inoculation. XJ44 was subsequently passaged in rhesus macaque lung fibroblasts *in vitro* and acquired an additional substitution in the transmembrane domain (TMD) of GP2^26^. This Phe to Ile mutation (F427I) has been found to be critical for attenuation in the mouse model ^26^. Introduction of this single substitution alone into the wild-type JUNV sequence significantly attenuates the virus in the suckling mouse model of JUNV lethal infection, as 90% of mice survived after intracranial inoculation. However, neither the glycan removal mutations nor the TMD substitution fully attenuate the virus, indicating that multiple mutations are necessary for complete attenuation.

Our previous work has demonstrated that a MACV strain expressing the whole GPC of Cd#1 is fully attenuated^27^. Likewise, a MACV strain containing the Cd#1 GPC ectodomain is also partially attenuated, though reversions at two N-linked glycans were frequently observed^28, 29^. Removal of the two N-linked glycans at N83 and N166 (MACV GPC_ΔN83/ΔN166_) results in a strain that is partially attenuated in IFN-αβ/γ R^−/−^ mice^29^. While 100% of these mice survived infection, they all developed non-lethal neurological signs of disease. Meanwhile, a MACV strain containing the analogous F438I TMD substitution was likewise partially attenuated in IFN-αβ/γ R^−/−^ mice^30^. However, 28% of mice infected with the MACV GPC_F438I_ mutants succumbed to disease and all viruses isolated reverted to wild type sequence.

In this study, we sought to fully attenuate MACV by combining mutations disrupting specific glycosylation sites with the TMD substitution. Herein, we report that a MACV mutant with glycan deficiencies at residues N83 and N166 and the F438I TMD substitution on GPC fully attenuates MACV in two different animal models. Furthermore, inoculation with this attenuated strain provides protection from subsequent lethal challenge with wild-type MACV. This protection can likely be attributed to a strong MACV-specific humoral response. Finally, we provide evidence that the F438I substitution leads to a replication defect in neurons, likely contributing to the attenuation phenotype observed.

## Results

### Introducing mutations to specific GPC glycan sites and the TMD F438I mutation leads to complete attenuation of MACV

Our previous work demonstrates that disruption of glycosylation sites at GPC N83 and N166 partially attenuate MACV in IFN-αβ/γ R^−/−^ mice^29^. Though all mice survive challenge with this mutant, all mice still develop neurological signs of disease. On the other hand, while the F438I substitution is known to abrogate neurological disease, it is genetically unstable *in vivo*^30^. We therefore combined the glycan removal mutations with the F438I substitution with the hypothesis that the combination of these mutations would lead to a more stable and complete attenuation. Using our established MACV reverse genetics system^31^, we created MACV mutants containing the F438I substitution in addition to glycan deficiency at either one or two sites (MACV GPC_ΔN83/F438I_, MACV GPC_ΔN166/F438I_, and MACV GPC_ΔN83/ΔN166/F438I_) (Fig. 1A). As 8.33% of the MACV GPC_ΔN83/F438I_ population in the passage 0 stock was found to be GPC_A83N_ and GPC_A85S_ revertants, we excluded MACV GPC_ΔN83/F438I_ in further studies. For each of the rescued viruses, GPC protein expression and cleavage was confirmed via Western blot (Fig. 1B). All these viruses displayed similar *in vitro* growth kinetics in Vero cells (Fig. 1C). Furthermore, no reversions or unintended mutations were found in the GPC gene after 5 serial passages in Vero cells (Supplementary Table 1).

**Figure 1.**
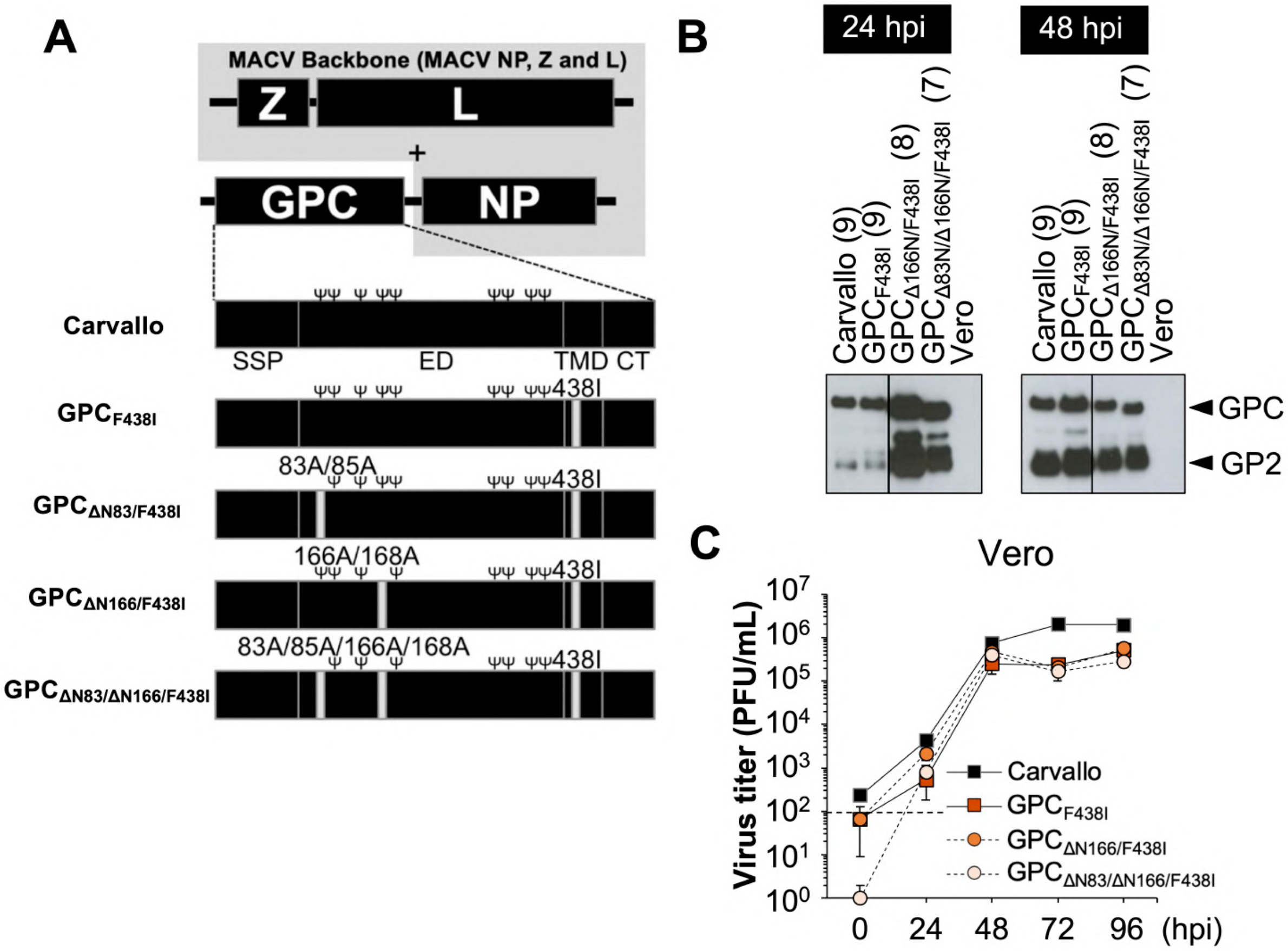
Creation of MACV with glycan removals and transmembrane substitution. (A) Schematic representation of the genome of MACVs. Within GPC, SSP (stable signal peptide), ED (ectodomain), TMD (transmembrane domain) and CT (cytoplasmic tail) are shown. Ψ in black character represents N-glycosylation site. (B) The representative WB of GPC in infected Vero cells at MOI = 1. The number of putative N-glycosylation sites is shown in parentheses. (C) Viral replication of MACV GPC_F438I_, MACV GPC_ΔN166/F438I_ and MACV GPC_ΔN83/ΔN166/F438I_ was characterized in Vero cells (MOI = 0.01). The dashed line indicates the detection limit. Data shown are averages of triplicate wells with error bars indicating the SD.

We next studied virus infection in the IFN-αβ/γ R^−/−^ mouse model. Groups of IFN-αβ/γ R^−/−^ mice were inoculated with MACV GPC_F438I_, MACV GPC_ΔN166/F438I_, MACV GPC_ΔN83/ΔN166/F438I_ or wild-type MACV Carvallo strain (originally isolated from a human patient’s spleen after a fatal case of BHF) and monitored for signs of disease for up to 49 days. Compared to MACV GPC_F438I_, removal of one or two glycans further attenuates the virus, with two glycan removals necessary for the most complete attenuation. 100% of mice infected with MACV GPC_ΔN83/ΔN166/F438I_ survived infection (Fig. 2A), and none of these mice developed any observable clinical illness (Fig. 2B). Furthermore, mice infected with MACV GPC_ΔN83/ΔN166/F438I_ maintained their body weights throughout the course of the study in contrast to animals infected with wild-type MACV (Fig. 2C). At time of death/euthanasia or at 49 days post-infection, serum, lung, liver, and brain samples were collected for viral titration and viral RNA detection. No virus was detected in any of the mice infected with MACV GPC_ΔN83/ΔN166/F438I_, while virus was detected in at least one animal from every other group (Fig. 2D). Likewise, viral RNA was not detected in the brains of MACV GPC_ΔN83/ΔN166/F438I_-infected animals (Fig. 2E and Supplementary Figure 1), further providing evidence that MACV GPC_ΔN83/ΔN166/F438I_ is highly attenuated in IFN-αβ/γ R^−/−^ mice.

**Figure 2.**
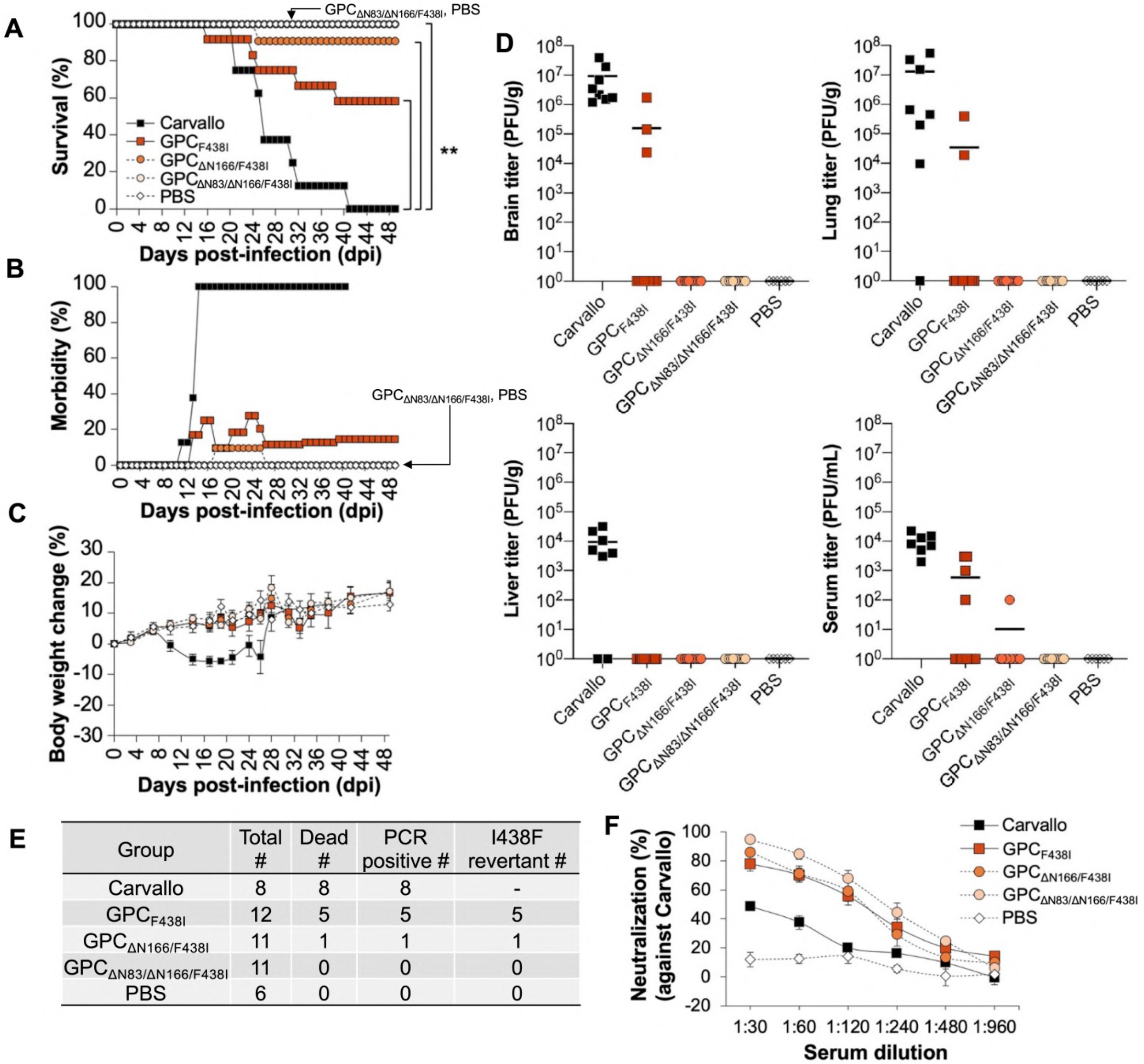
MACV with glycan removals and transmembrane substitution is attenuated in IFN-αβ/γ R^−/−^ mice. (A) Survival rate of IFN-αβ/γ R^−/−^mice after intraperitoneal injection of 10^4^ PFU of the indicated strains and mutants (Carvallo N=8, MACV GPC_F438I_ N=12, MACV GPC_ΔN166/F438I_ N= 11, MACV GPC_ΔN83/ΔN166/F438I_ N= 11, PBS N=6). Statistically significant differences are indicated by asterisks (**, P<0.01 by log rank test). (B) Morbidity rate of the IFN-αβ/γ R^−/−^ mice after infection. (C) Body weight changes were monitored on the indicated days. Error bars indicate the SEM. The data shown are pooled from two independent experiments. (D) Virus titers in brain, lung, liver and serum. The minimum detection limit is shown as a dashed line. Data under detection limit is plotted as “1 PFU/g or mL”. The solid bar represents the mean of the titers. (E) Viral RNA at time of euthanasia. Shown are the total number of animals per group, the number of animals per group that succumbed to infection and were humanely euthanized, the number of animals with detectable viral RNA (GPC) at time of death, and the number of animals with I438F revertant viral RNA. (F) Neutralization activity of mouse serum against wild-type MACV (Carvallo).

To determine if MACV GPC_ΔN83/ΔN166/F438I_ induces a neutralizing antibody response, we performed neutralizing assays for serum samples collected at 49 dpi (Fig 2F). MACV GPC_ΔN83/ΔN166/F438I_ elicited the strongest neutralizing antibody response against wild-type MACV Carvallo when compared to the other viruses tested. Our PRNT_50_ data also suggested that loss of N-linked glycans might enhance immunogenicity (Supplementary Table 2). To evaluate the effect of the N-linked glycans at N83 and N166 on viral sensitivity to antibody neutralization, we tested sera from MACV GPC_F438I_-infected animals against MACV GPC_ΔN166/F438I_ and MACV GPC_ΔN83/ΔN166/F438I_. The serum samples showed strong activity against MACV mutant with single glycan-deficiency (>217.4 for MACV GPC_ΔN166/F438I_) but had much stronger activity against MACV GPC_ΔN83/ΔN166/F438I_ (>646), which lacks both of these two N-glycans. These results indicate that the number of glycans on GPC is inversely correlated with the susceptibility of MACV to neutralizing antibodies, likely due to glycan shielding of neutralizing epitopes^32, 33^.

### MACV GPC_ΔN83/ΔN166/F438I_ is attenuated in outbred guinea pigs

MACV GPC_ΔN83/ΔN166/F438I_ was fully attenuated even after infection of immunocompromised mice, which are normally highly susceptible to the virus. Therefore, we sought to test whether this mutant would work as a vaccine in an immunocompetent guinea pig model. Before that, we first infected animals with the MACV GPC_ΔN83/ΔN166/F438I_ to confirm that this mutant virus does not cause disease in the guinea pigs. The Carvallo strain of MACV used in our previous mouse experiments is not fully lethal in the Hartley guinea pig model^34^. Therefore, to test the attenuation of our experimental vaccine we compared it to the lethal Chicava strain of MACV in Hartley guinea pigs^35^. Groups of guinea pigs were challenged intraperitoneally with 10,000 PFU of either MACV GPC_ΔN83/ΔN166/F438I_ or wild-type MACV (Chicava) and monitored for disease signs. Weights and temperatures were collected daily for the first 30 days post-infection. Consistent with our mouse experiments, 100% of guinea pigs infected with MACV GPC_ΔN83/ΔN166/F438I_ survived challenge, while 100% of animals infected with wild-type MACV succumbed to challenge by day 22 (Fig. 3A). Guinea pigs infected with wild-type MACV developed mild fevers as early as day 5 post-infection, and began to lose weight at day 15 post-infection, reaching humane euthanasia criteria by days 19-22. Some animals also developed additional clinical signs, including vomiting and hind limb paralysis, with the latter a clear manifestation of neurological pathology. By contrast, none of the animals infected with MACV GPC_ΔN83/ΔN166/F438I_ developed any clinical signs (Fig. 3B). These animals steadily gained weight over the course of the study (Fig. 3C), and none of the animals developed an elevated temperature (Fig. 3D).

**Figure 3.**
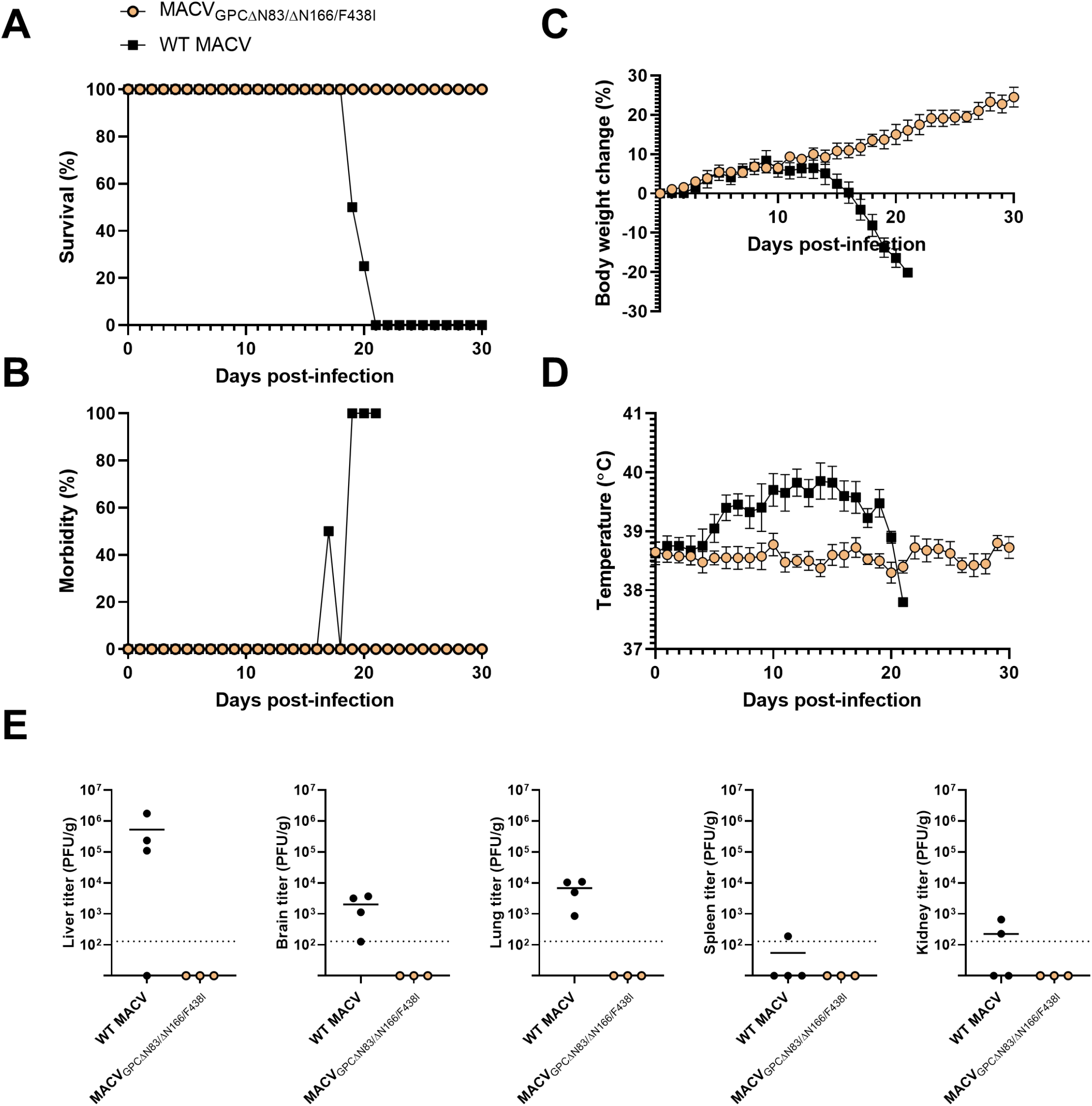
MACV with glycan removals and transmembrane substitution is attenuated in Hartley guinea pigs. (A) Survival rate of animals injected intraperitoneally with 10^4^ PFU of either MACV GPC_ΔN83/ΔN166/F438I_ (N = 10) or WT MACV (N = 4). Data is representative of two independent experiments. (B) Morbidity rate of animals after infection. Data is representative of two independent experiments. (C) Body weight changes were monitored daily for the first 30 days post-infection (N = 4 per group). Error bars represent the SEM. (D) Temperature was monitored via BMDS transponder daily for 30 days post-infection (MACV GPC_ΔN83/ΔN166/F438I_ N=4, WT MACV N=8). Error bars represent the SEM. (E) Virus titers in brain, liver, lung, spleen, and kidney. The minimum detection limit is shown as a dashed line. Data under detection limit is plotted as “1 PFU/g or mL”. The solid bar represents the mean of the titers.

Though no clinical illness was observed for any animal infected with MACV GPC_ΔN83/ΔN166/F438I_, we wanted to determine whether live virus and/or viral RNA was present in any organs. At 30 days post-infection, 3 animals infected with MACV GPC_ΔN83/ΔN166/F438I_ were humanely euthanized, and blood and organs were collected. We could not detect live virus in the brain, liver, spleen, lung, or kidney of any animal infected with MACV GPC_ΔN83/ΔN166/F438I_ (Fig. 3E). Furthermore, no viral RNA could be detected in any organ (data not shown). By contrast, animals infected with wild-type MACV all had detectable virus loads in at least one of the five organs tested at time of euthanasia. While some of these animals were able to clear the virus in some organs, all animals infected with wild-type MACV had detectable live virus in the brain. In summary, these results demonstrate that MACV GPC_ΔN83/ΔN166/F438I_ (Carvallo backbone) does not cause detectable disease in an immunocompetent animal model.

### Immunization with experimental vaccine MACV GPC_ΔN83/ΔN166/F438I_ protects guinea pigs from heterologous lethal challenge with MACV

Since MACV GPC_ΔN83/ΔN166/F438I_ is fully attenuated in both immunocompromised mice and immunocompetent guinea pigs, we next investigated whether this mutant MACV could protect animals from heterologous lethal challenge with wild-type MACV. In the MACV GPC_ΔN83/ΔN166/F438I_ group, guinea pigs were first immunized with MACV GPC_ΔN83/ΔN166/F438I_ and then challenged with 10,000 PFU of wild-type MACV (Chicava strain) at 60 days later. As a control, a new group of age-matched, naïve guinea pigs was challenged with 10,000 PFU of wild-type MACV (the control group). Animals were monitored for clinical signs of disease, and weight and temperature changes for 30 days post-challenge. Again, 100% of naïve control animals succumbed to wild-type challenge between days 19-22, with similar disease development as what was observed and reported previously. All animals immunized with MACV GPC_ΔN83/ΔN166/F438I_ survived challenge (Fig. 4A) without discernible clinical signs observed in any of the immunized animals (Fig. 4B). These animals maintained their weight throughout the course of the study (Fig. 4C) and never developed fevers (Fig. 4D), in contrast to the unimmunized, control animals.

**Figure 4.**
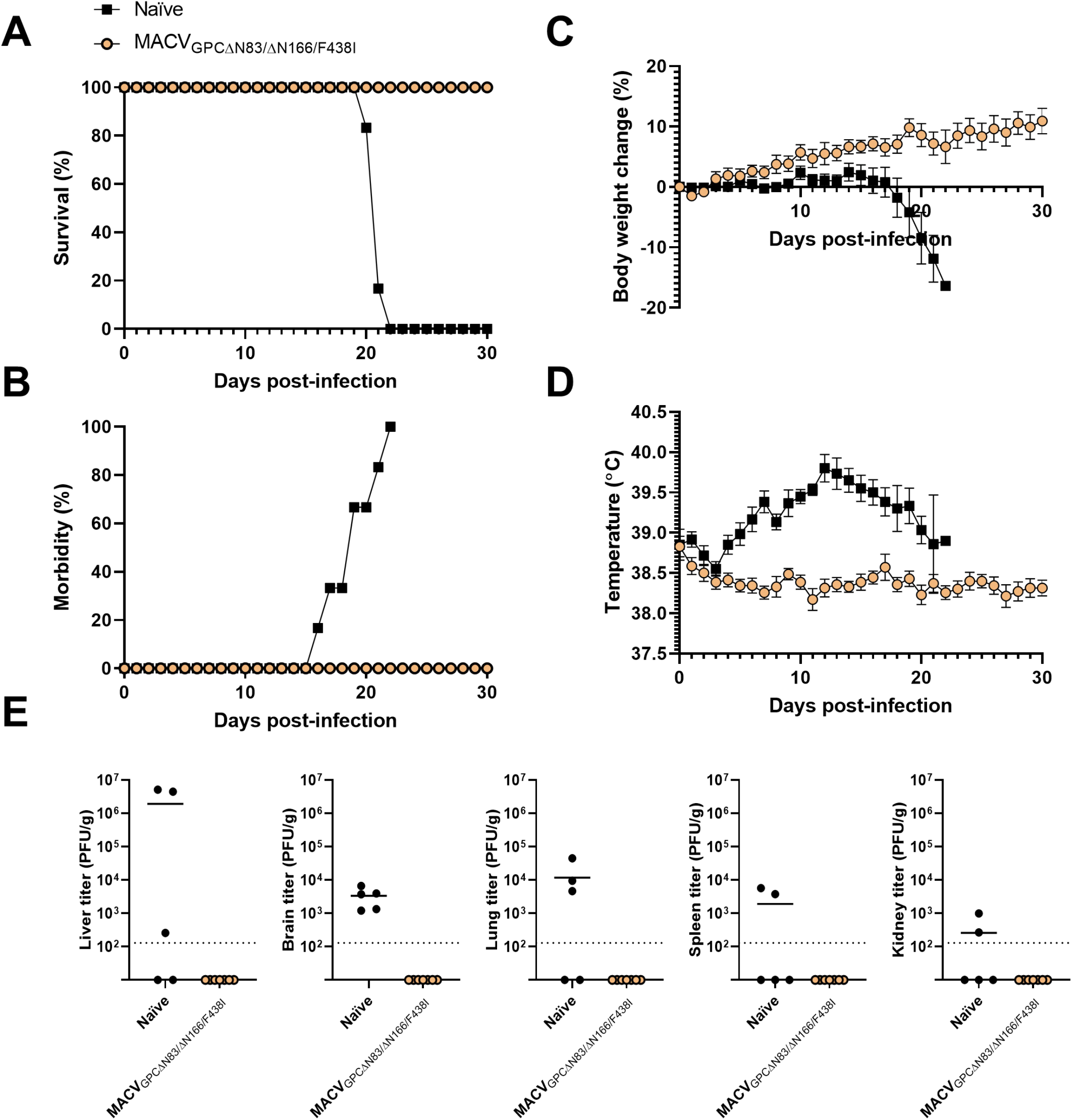
Guinea pigs challenged with wild-type MACV after inoculation with MACV GPC_ΔN83/ΔN166/F438I_. (A) Survival rate of animals inoculated with MACV GPC_ΔN83/ΔN166/F438I_ (N = 7) or naïve animals (N=6) challenged 60 days post-inoculation with an intraperitoneal injection of 10^4^ PFU of wild-type MACV. Data is representative of two independent experiments. (B) Morbidity rate of animals after infection. Data is representative of two independent experiments. (C) Body weight changes were monitored daily for the first 30 days post-infection. Error bars represent the SEM. Data is pooled from two independent experiments. (D) Temperature was monitored via BMDS transponder daily for 30 days post-infection. Error bars represent the SEM. Data is pooled from two independent experiments. (E) Virus titers in brain, liver, lung, spleen, and kidney. The minimum detection limit is shown as a dashed line. Data under detection limit is plotted as “1 PFU/g or mL”. The solid bar represents the mean of the titers.

30 days after the wild-type MACV challenge, animals inoculated with MACV GPC_ΔN83/ΔN166/F438I_ were humanely euthanized for the collection of blood and tissue samples. No live virus was detected in the brain, liver, spleen, lung, or kidney of any of the animals in the MACV GPC_ΔN83/ΔN166/F438I_ group (Fig. 4E). Consistently, viral RNA was undetectable (data not shown). By contrast, virus could be detected in brains of all animals in the control group. High viral titers were observed in other organs in the control group as well.

We performed a histopathological analysis of the liver, spleen, and brain of MACV-challenged Hartley guinea pigs. Portal and lobular inflammatory lymphohistiocytic lesions were observed in the livers of lethally challenged animals without immunization (Fig. 5, left panel). Mildly decreased white pulp cellularity was observed in the spleens of succumbed control animals, but not in those inoculated with MACV GPC_ΔN83/ΔN166/F438I_. Perivascular cellular infiltrates were observed in the brains of the control animals, but not in those inoculated with MACV GPC_ΔN83/ΔN166/F438I_ (Fig. 5)^35^. Overall, these results demonstrate that MACV GPC_ΔN83/ΔN166/F438I_ administration conferred protection from lethal MACV challenge.

**Figure 5.**
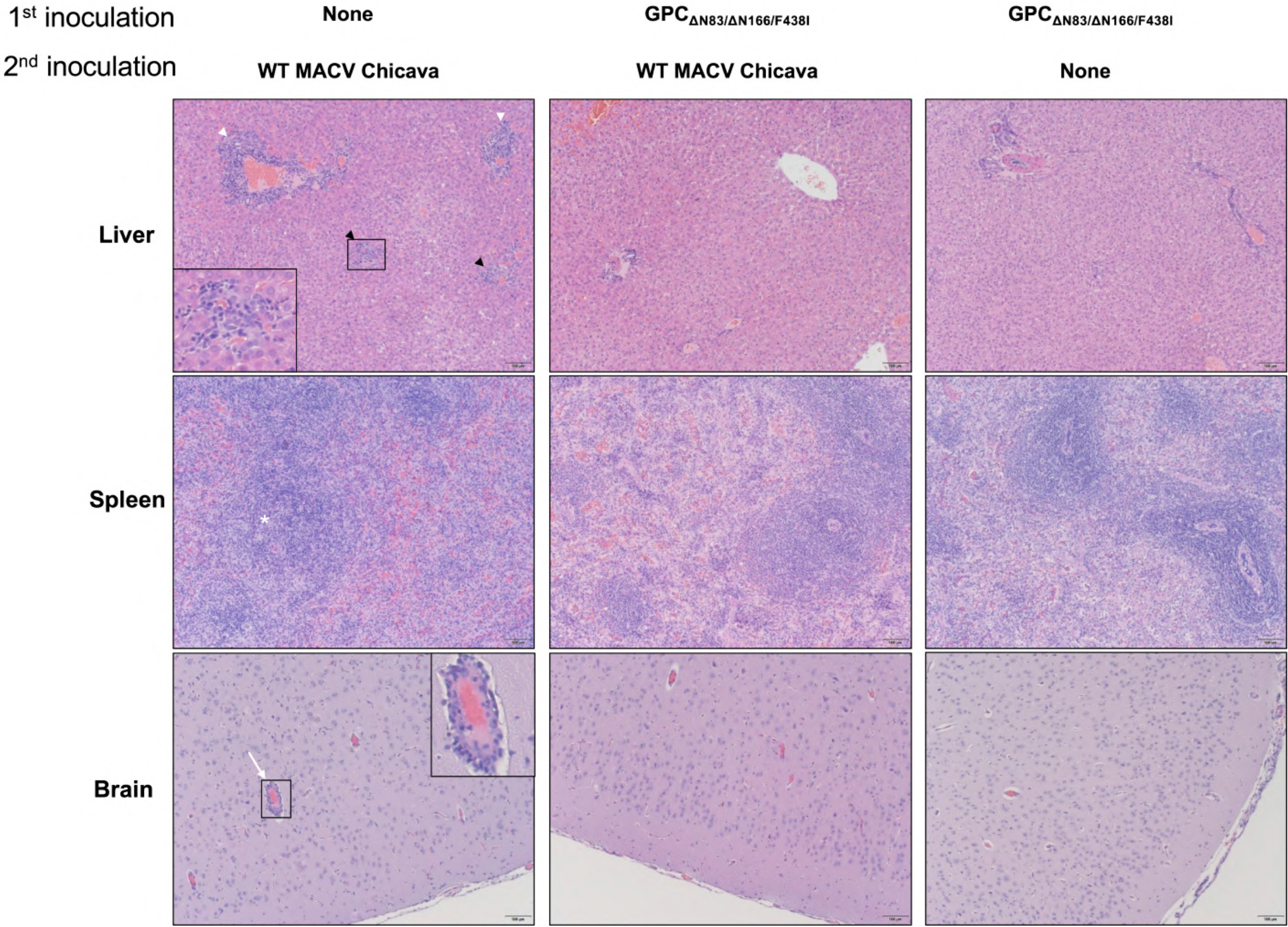
Histopathology from MACV-challenged Hartley guinea pigs. Major target organs of MACV were collected at time of euthanasia, 20-22 days after inoculation with Chicava without immunization (left panels), 90 days after inoculation with MACV GPC_N83/N166/F438I_ and 30 days following wild-type Chicava challenge (middle panels) or 30 days after inoculation with MACV GPC_N83/N166/F438I_ followed by no challenge (right panels). Liver: Portal (white arrowhead) and lobular (black arrowhead) inflammatory lymphohistiocytic lesions were observed in lethally challenged animals (inset: lobular focus of inflammation). Spleen: Mildly decreased white pulp cellularity (white asterisk) was observed in lethally challenged animals. Brain: Perivascular infiltrates were observed in lethally challenged animals (white arrow and inset). Hematoxylin and eosin stain, Magnification ×10.

### MACV-specific humoral response is generated after immunization with MACV GPC_ΔN83/ΔN166/F438I_

The humoral immune response is known to be critical for combating New World arenavirus infections^36, 37^, so we next wanted to determine whether MACV GPC_ΔN83/ΔN166/F438I_ elicited a humoral response that mediates the protection. Blood was taken from 3 animals at 30 days post-inoculation with MACV GPC_ΔN83/ΔN166/F438I_. Blood was also collected at 60 days post-inoculation from 4 animals inoculated with MACV GPC_ΔN83/ΔN166/F438I_ as well as 3 naïve control animals. We first performed PRNT assays to assess neutralizing antibody titers against wild type MACV (both Carvallo and Chicava strain) as well as the homologous MACV GPC_ΔN83/ΔN166/F438I_ mutant. Six out of seven animals developed a strong neutralizing antibody response to the glycan deficient MACV GPC_ΔN83/ΔN166/F438I,_ which increased over time. However, six of seven animals did not develop neutralizing antibodies against either wild-type MACV strain (Fig. 6A). This is consistent with previous data demonstrating that MACV mutants lacking specific glycans on GPC are more susceptible to antibody neutralization^29^. We further performed ELISAs to measure the level of MACV GP1 and NP specific antibodies with purified MACV (Carvallo) GP1 and NP proteins. All but one animal had detectable MACV NP or GP1 binding antibodies, though titers were highly variable amongst individual animals (Fig. 6B). Overall, these results demonstrate that most of the animals inoculated with the glycan deficient MACV GPC_ΔN83/ΔN166/F438I_ developed a measurable humoral response; however, the neutralizing antibody response to wild type MACV was below the detection limit.

**Figure 6.**
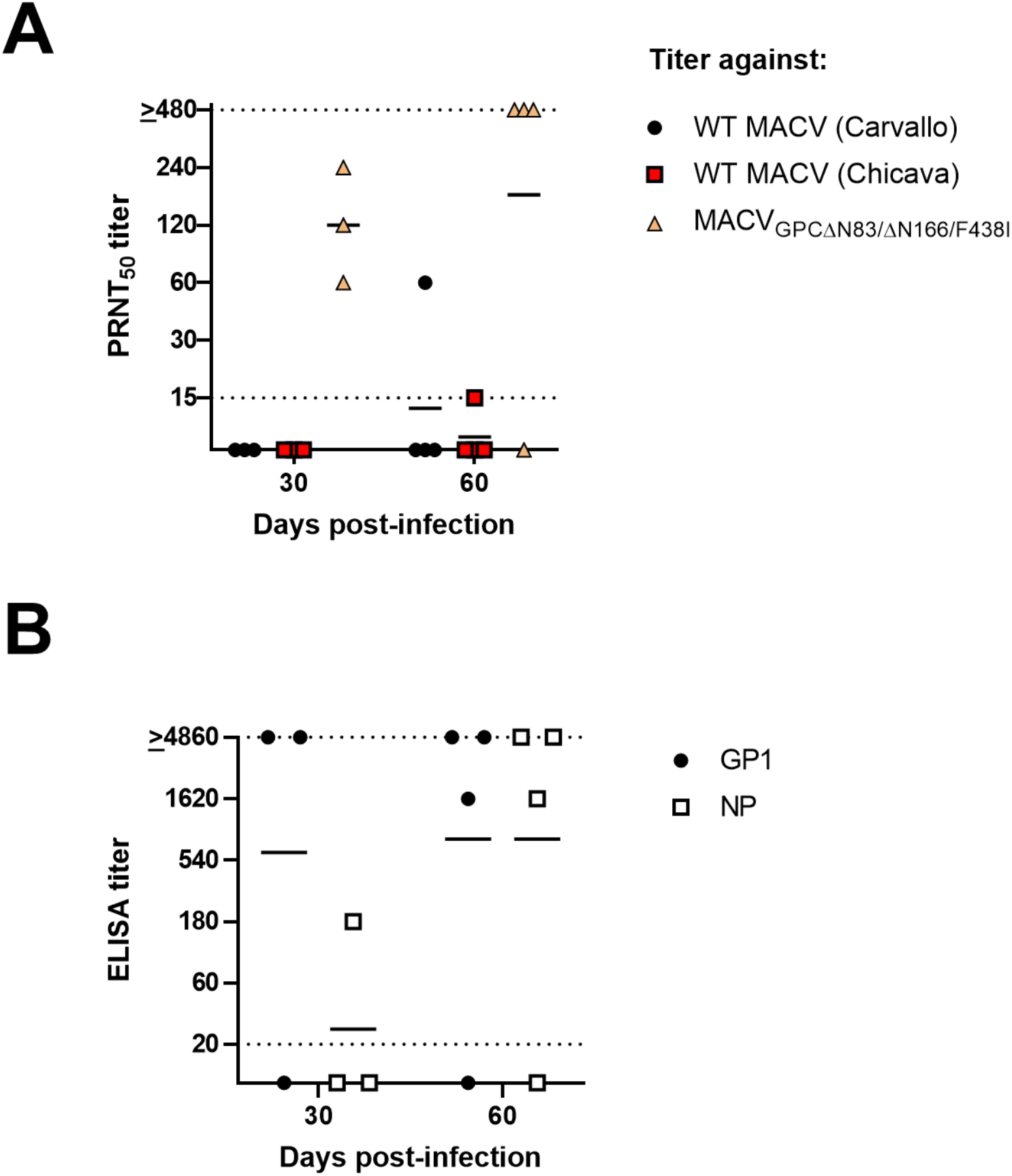
Humoral response following inoculation with attenuated MACV. (A) PRNT_50_ titers of guinea pig serum collected at either 30 days or 60 days post-inoculation with MACV GPC_ΔN83/ΔN166/F438I._ Neutralizing activity against Carvallo, Chicava, and GPC_ΔN83/ΔN166/F438I_ mutants of MACV was assessed. The solid line represents the geometric mean, while the dashed lines indicate the upper and lower limits of detection. (B) MACV-specific antibodies against either GP1 or NP measured via ELISA after 30 or 60 days post-inoculation with MACV GPC_ΔN83/ΔN166/F438I._ An ELISA titer was considered positive when the OD_450_ value was higher than at least two standard deviations of the mean of naïve animal serum run concurrently.

### MACV containing F438I substitution in GPC TMD replicates poorly in neurons

It is clear that the F438I substitution in MACV GPC abrogates neurological pathology in our IFN-αβ/γ R^−/−^ mouse model. However, the mechanism by which the F438I substitution in GPC TMD attenuates MACV *in vivo* remains unknown. To address this question, we compared the growth of wild-type MACV (Carvallo), MACV GPC_F438I_, MACV GPC_ΔN166/F438I_ and MACV GPC_ΔN83/ΔN166/F438I_ in mouse astrocytoma (C8D1A) and neuroblastoma (Neuro 2a) cell lines at an MOI of 0.01. While we observed no significant growth differences for these viruses in Vero cells (Fig. 1C) and mouse astrocytes (C8D1A) (Fig. 7A), all mutants containing the F438I substitution replicated to significantly lower titers than wild-type MACV in mouse neurons (Neuro 2a) (Fig. 7A). MACV GPC_F438I_ demonstrated the most severe defect in virus growth, while removal of glycans appeared to partially restore replication. We further confirmed this observation in human astrocytoma (132N1) and neuroblastoma (IMR-32) cell lines. Again, we observed no growth differences in 132N1 astrocytes, but we detected a significantly impaired growth of all mutant strains in human IMR-32 neurons (Fig. 7B). These data demonstrated for the first time that the F438I substitution affects MACV infection in cultured neurons, providing one potential mechanism for the attenuation *in vivo*.

**Figure 7.**
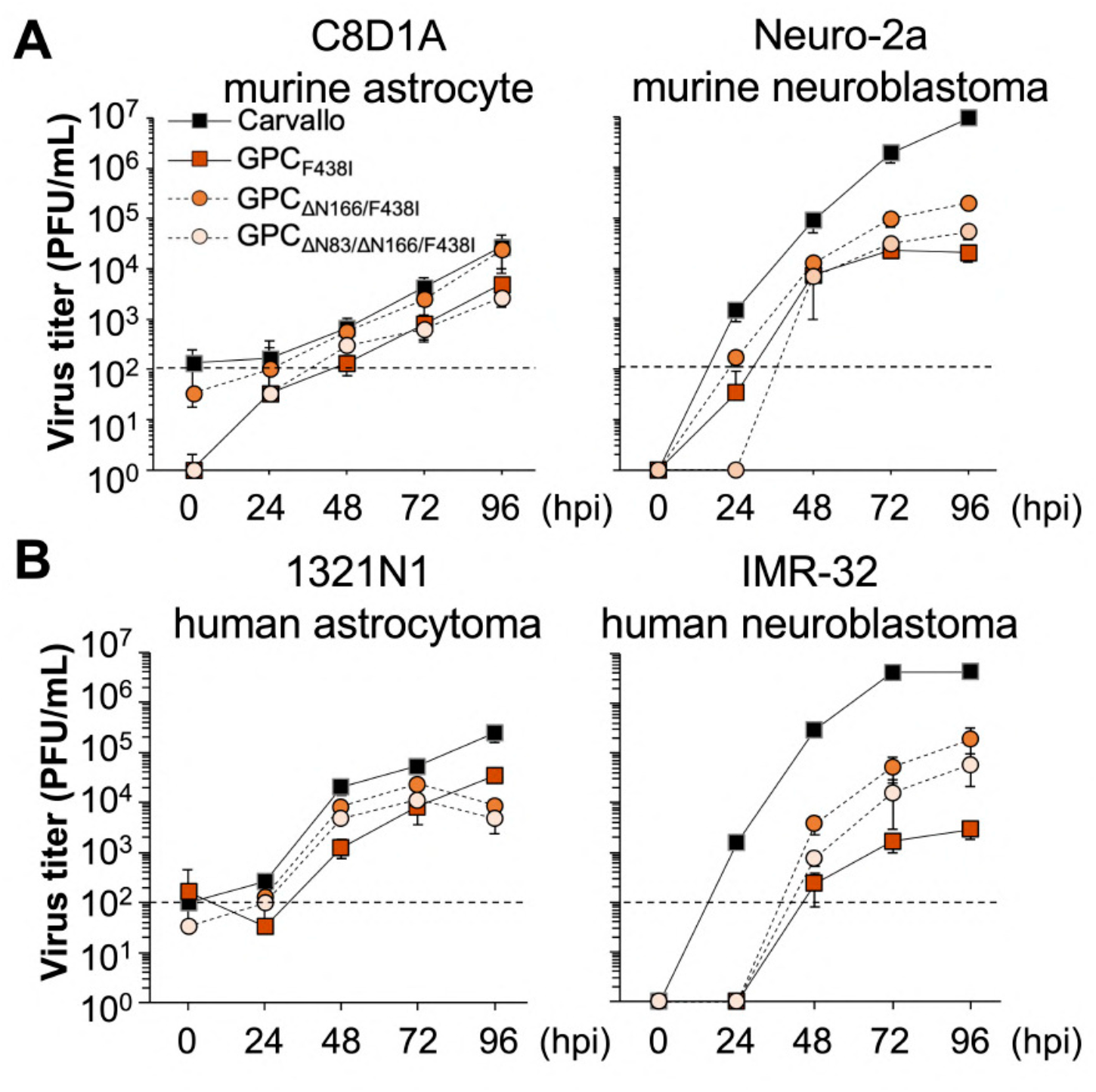
Replication of MACV containing the F438I transmembrane substitution in neurological cell lines. (A) Viral replication of MACV GPC_F438I_, MACV GPC_ΔN166/F438I_ and MACV GPC_ΔN83/ΔN166/F438I_ was characterized in C8D1A and Neuro-2a cells (MOI = 0.01). The dashed line indicates the detection limit. Data shown are averages of triplicate wells with error bars indicating the SD. (B) Viral replication of MACV GPC_F438I_, MACV GPC_ΔN166/F438I_ and MACV GPC_ΔN83/ΔN166/F438I_ was characterized in human-derived cell lines 1321N1 astrocytes and IMR-32 neurons (MOI = 0.01). The dashed line indicates the detection limit. Data shown are averages of triplicate wells with error bars indicating the SD.

## Discussion

In this study, we characterized MACV mutant containing the TMD F438I substitution and mutations at glycosylation sites on GP1. Our previous research revealed that neither the F438I substitution nor the glycosylation site mutations alone are sufficient for full attenuation in IFN-αβ/γ R^−/−^ mice^29, 30^. This work demonstrates that the combination of the F438I substitution in TMD with the glycan deficiency at N83 and N166 of GPC led to full attenuation in our IFN-αβ/γ R^−/−^ mouse model. Interestingly, both glycans must be absent for the complete and stable attenuation. Lack of the glycan at N166, even along with the F438I mutation, is not sufficient for full attenuation in mice, with one out of 10 mice developing signs of disease and ultimately succumbing to challenge. Sequencing confirmed that the GPC of the MACV from the one mouse that succumbed to disease contained the I438F reversion, which is likely the cause of the observed disease in this animal. We did not observe any reversion of MACV GPC_ΔN83/ΔN166/F438I_ *in vitro*, either in Vero or Neuro-2a cells (Supplementary Table 1). However, we could not isolate live virus from any animal infected with MACV GPC_ΔN83/ΔN166/F438I_, so it is unclear whether reversion occurred *in vivo*. Further studies attempting to isolate live virus from animals earlier in the course of infection will be needed to assess the genetic stability of MACV GPC_ΔN83/ΔN166/F438I_ *in vivo*.

We did not conclusively determine that MACV-specific antibodies play a significant role in mediating either the attenuation of MACV GPC_ΔN83/ΔN166/F438I_ or the protection from lethal challenge observed during this study. We did observe that guinea pigs inoculated with MACV GPC_ΔN83/ΔN166/F438I_ developed high titers of neutralizing antibodies against MACV GPC_ΔN83/ΔN166/F438I_. However, interestingly, none of the animals developed detectable neutralizing antibodies against either the wild-type Carvallo or Chicava strain of MACV by day 30 post-infection, and only one animal had developed low titers of neutralizing antibodies against the wild-type strains by day 60 post-infection. Previous studies have determined that neutralizing antibodies against JUNV do not neutralize MACV due to a loop 10 feature that blocks the receptor binding domain (RBD)^38,39^. Glycans likely play a similar role in blocking the RBD from neutralization, as we observe that MACV GPC_ΔN83/ΔN166/F438I_ is more easily neutralized by antibodies due to the loss of glycans, and that the wild-type viruses are less easily neutralized likely due to glycan shielding^32, 33, 40^. Nevertheless, despite the lack of detectable neutralizing antibodies against wild-type MACV, all of our guinea pigs survived subsequent wild-type challenge with no clinical signs noted, indicating that protection is not mediated by neutralizing antibodies alone. All except one animal developed high MACV-specific antibodies, perhaps indicating that antibody-dependent cellular cytotoxicity (ADCC) and other non-neutralizing antibody functions contribute to protection. Non-neutralizing antibodies against the GP of ebolavirus and the HA of influenza have been demonstrated to offer protection^41–43^. The role of non-neutralizing antibodies during arenavirus infection has also been noted as significant in several studies. Non-neutralizing, binding antibodies directed against LCMV have been demonstrated to accelerate viral clearance from infected mice^44, 45^. For LASV, a vaccine candidate consisting of vaccinia virus containing LASV NP RNA provided some protection despite the absence of GPC^46^. In addition, an inactivated rabies vaccine vector expressing LASV GPC failed to elicit a neutralizing antibody response in vaccinated mice yet did elicit high titers of non-neutralizing antibodies capable of stimulating ADCC^47^. Another study demonstrated that, while full-length IgG antibodies that neutralize JUNV *in vitro* protect guinea pigs from lethal challenge with JUNV, F(ab’)2 fragment antibodies that still successfully neutralize JUNV *in vitro* do not offer protection for guinea pigs infected with JUNV^48^. This indicates that elimination of infected cells via Fc-mediated effector functions is critical. For MACV-specific antibodies, ADCC activity has been observed^14^, but further studies will be needed to elucidate the role of non-neutralizing antibodies during New World arenavirus infection.

Cellular immunity may also play a role, as one of our animals had no detectable MACV-specific antibodies of any type but nevertheless survived wild-type challenge. This animal later developed MACV-specific antibodies after the wild-type challenge (Supplementary Figure 2), confirming that the animal was infected and yet survived challenge. Because these animals were outbred, variability in their response to MACV infection is expected. Only one-third of human BHF patients develop severe disease^5^, so the impact of other host factors cannot be ruled out. Nevertheless, further studies examining the potential contributions of cellular immunity during MACV infection would be beneficial.

Another important observation from our study was the important contribution of the F438I substitution to the attenuation of the virus in the nervous system. In our previous study, 100% of MACV GPC_ΔN83/ΔN166_-infected IFN-αβ/γ R^−/−^ mice developed neurological signs of disease such as imbalance. In this study, addition of the F438I substitution produced a mutant incapable of causing observable neurological signs of disease in both IFN-αβ/γ R^−/−^ mice and Hartley guinea pigs. For both of our animal models, 100% of naïve animals infected with wild-type MACV developed detectable viral titers in the brain, even late in infection after the virus has been cleared from other organs. Meanwhile, no live virus or viral RNA could be detected in the brains of any animal inoculated with MACV GPC_ΔN83/ΔN166/F438I_. We cannot confirm in our animal models whether MACV GPC_ΔN83/ΔN166/F438I_ can invade the nervous system but is incapable of replicating in the brain or whether it is incapable of spreading to the nervous system altogether. Future studies using an intracranial mode of inoculation would better address this issue. However, we did confirm that MACV GPC_ΔN83/ΔN166/F438I_ replicates poorly relative to wild-type virus in both mouse and human neurons *in vitro*. In our study, the combination of the glycan removals and the F438I substitution led to an overall significant loss of replication in neurons, which may partially explain the attenuation observed.

The mechanism behind the replication defect of MACV GPC_ΔN83/ΔN166/F438I_ and MACV GPC_F438I_ in neurons remains unclear. One previous report indicated that the F427I substitution in JUNV promotes cell-to-cell fusion at a neutral pH, as opposed to the acidic pH normally required for fusion, which led to reduced JUNV minigenome replication^49^. The authors speculated that the addition of the F427I substitution alters the GP in such a way that it remains in a post-fusion conformation. However, other studies have reported that the pH needed for fusion remains unchanged in JUNV mutants containing F427I alone^50^. Nevertheless, these reports also demonstrate that a substitution at F427 can restore the phenotype of SSP mutants that lower the required pH for fusion^18, 50, 51^. The F427 residue has been demonstrated to genetically interact with certain charged residues in the SSP^50, 52^. In addition to a potential defect in entry/fusion, a potential conformational mismatch with SSP due to the F427I substitution could lead to a reduction in GPC protein trafficking as well. Interestingly, we observed a partial restoration in replication in MACV GPC_ΔN83/ΔN166/F438I_ compared to MACV GPC_F438I_. One previous report has indicated that glycan removal in the LCMV GPC also leads to enhanced replication in neurons^53^. This could be due to a loss of steric hindrance from the N166 site, allowing for easier access to the cellular receptor. Future studies should focus on determining the exact mechanism for attenuation of these viruses that is so specific to neurons.

Overall, the results from our study demonstrate that a combination of mutations at glycosylation sites and the TMD of the arenavirus GPC is necessary for the complete attenuation. We have demonstrated that the MACV GPC_ΔN83/ΔN166/F438I_ is attenuated in two animal models, a key requirement in the development of potential vaccine candidates. Furthermore, immunization with MACV GPC_ΔN83/ΔN166/F438I_ is sufficient to protect guinea pigs from a lethal dose of wild-type MACV. These results emphasize the necessity of including multiple attenuating mutations when designing attenuated arenavirus mutants. This is particularly important for the creation of potential vaccine candidates, as multiple mutations reduce the chance of genetic reversion and thus increase the safety of live-attenuated vaccines. Taken together, our results demonstrate the feasibility of this approach in the rational design of future arenavirus vaccine candidates. Furthermore, it might be possible to generate genetically stable mutants that are highly attenuated and safe for use in diagnostic, antiviral, and molecular research outside of high containment.

## Figure Legends

**Supplementary Figure 1. Genetic stability of MACV with F438I substitution**.

All PCR positive animals succumbed to infection and had Phe at GPC 438. M34 and M36 in Trial 2 were negative in the first PCR. However, because they were symptomatic mice, the sample amount was increased, and the second PCR was performed (see Trial 2 ^2nd^). M36 in Trial 2 was a survivor but showed a mild scruffy coat and hunched posture.

**Supplementary Figure 2. Humoral response following challenge with wild-type MACV**. (A) PRNT_50_ titers of guinea pig serum collected at either time of death or at 30 days post-challenge with wild-type MACV. Neutralizing activity against Carvallo, Chicava, and GPC_ΔN83/ΔN166/F438I_ strains of MACV was assessed. The solid line represents the geometric mean, while the dashed lines indicate the upper and lower limits of detection. (B) MACV-specific antibodies measured via ELISA at time of death or at 30 days post-challenge with wild-type MACV. An ELISA titer was considered positive when the OD_450_ value was higher than at least two standard deviations of the mean of naïve animal serum run concurrently.

**Supplementary Table 1**. GPC sequences after serial passage *in vitro*.

**Supplementary Table 2**. PRNT_50_ geometric mean titer in mice infected with MACV mutants

## Materials & Methods

### Cells and viruses

Vero cells (ATCC, CCL-81), Neuro-2a (ATCC, CCL-131), C8D1A (ATCC, CRL-2541), 1321N1 (Millipore Sigma), IMR-32 (ATCC, CCL-127) and BHK-21 cells (ATCC, CCL-10) were maintained in minimal essential medium (MEM) (GE Healthcare Life Sciences) supplemented with 10% FBS (Life Technologies) and 1% penicillin-streptomycin (P/S)(Life Technologies) at 37°C with 5% CO2. HEK-293T cells (Life Technologies) were maintained in Dulbecco’s modified Eagles medium (DMEM) (GE Healthcare Life Sciences) with 10% FBS (Life Technologies) and 1% P/S (Life Technologies) at 37°C with 5% CO_2_. The recombinant MACV Carvallo strain^31^, MACV GPC_F438I_^30^, MACV GPC_ΔN83/ΔN166_^29^, MACV GPC_ΔN83/F438I_, MACV GPC_ΔN166/F438I_, and MACV GPC_ΔN83/ΔN166/F438I_ were rescued by using a reverse genetics system previously described^31, 54^. The wild-type Chicava strain was a generous gift from Dr. Thomas Ksiazek. We used the first or second passage of viruses for all infection experiments in this study. For *in vitro* infection, confluent monolayers of Vero, C8D1A or Neuro-2a cells were infected with viruses at MOI=0.01 for virus growth curves or MOI=1 for protein expression profiling. After 1h incubation at 37°C with 5% CO_2_, medium was replaced with MEM containing 2% FBS and 1% P/S. The supernatants or cells were collected at the indicated days. Virus titers of serum samples, homogenized organs or supernatant from cell culture were determined by plaque assay with tragacanth gum as previously described^28^. All experiments with live viruses except Cd#1 were performed in the BSL-4 laboratories at the Galveston National Laboratory (GNL) in accordance with institutional safety guidelines, NIH guidelines and US federal law.

### Animal studies

IFN-αβ/γ R^−/−^ mice were bred and maintained in the ABSL-2 facilities in the GNL at the University of Texas Medical Branch at Galveston. Five to nine week-old IFN-αβ/γ R^−/−^ mice were challenged by intraperitoneal injection with recombinant MACVs (10,000 PFU) and monitored for 49 days post-infection (dpi). All animal experiments were performed twice in independent studies. Serum, brain and liver samples were collected for virus titration and virus RNA copies when animals were euthanized or dead. Animals were humanely euthanized at the end of study (49 dpi) or if they became paralyzed or lost more than 20% of body weight. The Institutional Animal Care and Use Committee at the University of Texas Medical Branch at Galveston approved the study protocol (1208050A).

Six to eight week-old Hartley guinea pigs (Charles River Laboratories) were challenged by intraperitoneal injection with 10,000 PFU of recombinant MACV GPC_ΔN83/ΔN166/F438I_ (Carvallo backbone) or wild-type MACV Chicava strain. Animals were monitored for up to 90 days post-infection. Weights were taken daily for the first 30 days post-infection. Temperatures were recorded daily for the first 30 days post-infection using a Bio Medic Data Systems (BMDS) transponder following subcutaneous insertion of a transponder chip in each animal. Blood was collected via the vena cava at 60 dpi and at time of euthanasia. At 60 dpi, surviving animals were re-challenged with 10,000 PFU of wild-type MACV (Chicava) intraperitoneally. Animals were humanely euthanized at 30 or 90 dpi or if they became paralyzed or lost more than 15% of their body weight. At time of euthanasia, brain, liver, spleen, lung, and kidney samples were collected for virus titration, viral RNA content, and histopathology. The Institutional Animal Care and Use Committee at the University of Texas Medical Branch at Galveston approved the study protocol (2006067).

### RNA extraction, cDNA synthesis and sequence analysis

RNAs were extracted from homogenized organs or lysed cell lines by using Trizol reagent (Life Technologies) and Direct-zol RNA MiniPrep kits (Zymo Research, Irvine, CA) as previously described^27^. Reverse transcription was performed by using the Superscript III First-Strand Synthesis System (Life Technologies) with random primers according to manufacturer’s protocol. cDNAs for GPC ORF were amplified by PCR with 5’-CGCACCGGGGATCCTAGGCGATTC-3’ and 5’-CCTCTCAGCCTTCTATTTCTACCC-3’. cDNAs for whole virus genome were amplified by PCR into two and three DNA fragments for viral S segment with previously described primers^28^ and L segments with the following primers, respectively: 5’-CGCACCGGGGATCCTAGGCGTAAC-3’ and 5’-TAGGAACTGTGCCAGAAAGG-3’, 5’-TCTCTAACGCACTTGCTACC-3’ and 5’-TTGGATGTGCTGTGGTGAAC-3’, 5’-GAGGATGTTGCCTAACTC-3’ and 5’-CGCACCGAGGATCCTAGGCGACAC-3’. To read the sequence of the 5’ and 3’ end, 5’ RACE System for Rapid Amplification of cDNA Ends (Life Technologies) and the 3’ RACE System for Rapid Amplification of cDNA Ends (Life Technologies) were used according to manufacturer’s protocol. After purification using a QIAquick PCR purification kit (Qiagen), PCR products were directly sequenced using an ABI Prism 3130xl DNA sequencer (Life Technologies).

### Western blotting

Western blotting was performed as previously described^22^. Briefly, infected or transfected cells were harvested in the 2x Laemmli Sample Buffer (Bio-Rad) containing 5% β-mercaptoethanol and boiled at 95°C for 5 min. The samples were loaded in the wells of 4-15% Mini-Protean TGX gels (Bio-Rad). Then, proteins were transferred to polyvinyl difluoride (PVDF) membranes using the Trans-Blot Turbo Transfer System (Bio-Rad). After blocking with PBS containing 0.05% Tween 20 (PBS-T) and 5% skim milk (blocking buffer) for 1h at room temperature, the membranes were reacted with primary antibodies in blocking buffer at 4°C overnight. After three washes with PBS-T, the membranes were reacted with secondary antibody in blocking buffer for 2 h at room temperature. After three washes with PBS-T, the HRP signal was visualized by enhanced chemoluminescence (ECL) (ECL Western Blotting System, Amersham). The primary polyclonal antibodies targeting the cytoplasmic tail of MACV GP2 were created by immunization of synthetic peptides (KYPRLKKPTIWHKR) (ProSci) in rabbit.

### Plaque reduction neutralization test

Serum samples were heat inactivated at 56°C for 30 min. The serum samples were diluted and mixed with an equal volume of diluent containing 80 PFU of each virus. After incubation for 1 h at 37°C, the mixture was applied on Vero cell monolayers. After an incubation for 1 h at 37°C, the inoculum was replaced with tragacanth overlay (1.2% tragacanth gum mixed with equal volume of Temin’s 2X MEM containing 4% FBS and 2% P/S) and incubated for 7 to 8 days. Then the plates were fixed and stained with 1% crystal violet in 10% formalin.

### Enzyme-linked immunosorbent assay (ELISA)

Purified MACV (Carvallo strain) GP1 or NP (Immune Technology) were used as antigens. ELISA plates were coated with 25 ng of antigen per well and incubated overnight at 4°C. Wells were then blocked with 5% skim milk suspended in PBS. After 1 wash with PBS-T, wells were incubated with serially diluted serum for 1 hour at room temperature. Wells were then washed 5 times with PBS-T before incubation with secondary antibody (1:10,000 diluted Goat Anti-Guinea pig IgG H&L (HRP) (Abcam) for 1 hour at room temperature. Following 5 more washes with PBS-T, 3,3’,5,5’-tetramethylbenzidine Liquid Substrate, Supersensitive, for ELISA (Sigma) was added to visualize the reaction. After 30 minutes of incubation at room temperature, 1M phosphoric acid was added to stop the reaction. Optical density at 450 nm was then measured. A positive titer was considered to be anything above two standard deviations of the value for the corresponding naïve control serum.

### Histopathological analysis

Tissues were collected from dead or euthanized animals and fixed in 10% buffered formalin for a minimum of 5 days. Tissues were then cut, paraffin-embedded, and sectioned into 5 μm slices for standard staining with hematoxylin and eosin (H&E).

### Statistical analysis

Data were analyzed using Dunnett’s post hoc test following a one-way analysis of variance (ANOVA), log rank analysis, and the Mann-Whitney U test. Results were considered to be statistically significantly different when the P value was <0.05.

## Acknowledgments

E.K.M was supported by National Institutes of Health T32 training grant AI060549. J.T.M received funding from the Clinical and Translational Science Award NRSA (TL1) Training Core (TL1TR001440) from the National Center for Advancing Translational Sciences at the National Institutes of Health. Work in the Paessler laboratory was supported in parts by Public Health Service grants RO1AI093445 and RO1AI129198 and the John. S. Dunn Distinguished Chair in Biodefense endowment. This work was performed in part to complete the dissertation requirements for E.K.M. We thank David H. Walker for consultation on histopathology data and Milagros Miller for her technical help with animals.

## References

1. Lassa Fever. Centers for Disease Control and Prevention; 2019.

2. Kerber R, Reindl S, Romanowski V, Gómez RM, Ogbaini-Emovon E, Günther S, ter Meulen J. Research efforts to control highly pathogenic arenaviruses: a summary of the progress and gaps. J Clin Virol. 2015;64:120–7. Epub 2014/12/18. doi: 10.1016/j.jcv.2014.12.004. PubMed PMID: 25549822.

3. Delgado S, Erickson BR, Agudo R, Blair PJ, Vallejo E, Albariño CG, Vargas J, Comer JA, Rollin PE, Ksiazek TG, Olson JG, Nichol ST. Chapare virus, a newly discovered arenavirus isolated from a fatal hemorrhagic fever case in Bolivia. PLoS Pathog. 2008;4(4):e1000047. Epub 2008/04/18. doi: 10.1371/journal.ppat.1000047. PubMed PMID: 18421377; PMCID: PMC2277458.

4. Enria DA, Maiztegui JI. Antiviral treatment of Argentine hemorrhagic fever. Antiviral Res. 1994;23(1):23–31. doi: 10.1016/0166-3542(94)90030-2. PubMed PMID: 8141590.

5. Patterson M, Grant A, Paessler S. Epidemiology and pathogenesis of Bolivian hemorrhagic fever. Curr Opin Virol. 2014;5:82–90. Epub 2014/03/15. doi: 10.1016/j.coviro.2014.02.007. PubMed PMID: 24636947; PMCID: PMC4028408.

6. Mackenzie RB. Epidemiology of Machupo virus infection. I. Pattern of human infection, San Joaquín, Bolivia, 1962-1964. Am J Trop Med Hyg. 1965;14(5):808–13. doi: 10.4269/ajtmh.1965.14.808. PubMed PMID: 5829142.

7. Peters CJ, Kuehne RW, Mercado RR, Le Bow RH, Spertzel RO, Webb PA. Hemorrhagic fever in Cochabamba, Bolivia, 1971. Am J Epidemiol. 1974;99(6):425–33. doi: 10.1093/oxfordjournals.aje.a121631. PubMed PMID: 4601699.

8. Grant A, Seregin A, Huang C, Kolokoltsova O, Brasier A, Peters C, Paessler S. Junín virus pathogenesis and virus replication. Viruses. 2012;4(10):2317–39. Epub 2012/10/19. doi: 10.3390/v4102317. PubMed PMID: 23202466; PMCID: PMC3497054.

9. M B, J dlT, C.P. Arenaviridae: The Viruses and Their Replication. In: Hp K, editor. Fields Virology. Philadelphia, PA, USA: Wolter Kluwer Lippincott Williams & Wilkins; 2007. p. 1791–827.

10. Oppliger J, da Palma JR, Burri DJ, Bergeron E, Khatib AM, Spiropoulou CF, Pasquato A, Kunz S. A molecular sensor to characterize arenavirus envelope glycoprotein cleavage by subtilisin kexin isozyme 1/site 1 protease. J Virol. 2016;90(2):705–14. Epub 2015/10/28. doi: 10.1128/JVI.01751-15. PubMed PMID: 26512085; PMCID: PMC4702697.

11. York J, Nunberg JH. Distinct requirements for signal peptidase processing and function in the stable signal peptide subunit of the Junín virus envelope glycoprotein. Virology. 2007;359(1):72–81. Epub 2006/10/12. doi: 10.1016/j.virol.2006.08.048. PubMed PMID: 17045626.

12. Lenz O, ter Meulen J, Klenk HD, Seidah NG, Garten W. The Lassa virus glycoprotein precursor GP-C is proteolytically processed by subtilase SKI-1/S1P. Proc Natl Acad Sci U S A. 2001;98(22):12701–5. Epub 2001/10/16. doi: 10.1073/pnas.221447598. PubMed PMID: 11606739; PMCID: PMC60117.

13. Burri DJ, da Palma JR, Kunz S, Pasquato A. Envelope glycoprotein of arenaviruses. Viruses. 2012;4(10):2162–81. Epub 2012/10/17. doi: 10.3390/v4102162. PubMed PMID: 23202458; PMCID: PMC3497046.

14. Amanat F, Duehr J, Huang C, Paessler S, Tan GS, Krammer F. Monoclonal antibodies with neutralizing activity and Fc-effector functions against the Machupo virus glycoprotein. J Virol. 2020;94(5). Epub 2020/02/14. doi: 10.1128/JVI.01741-19. PubMed PMID: 31801871; PMCID: PMC7022345.

15. Cao W, Henry MD, Borrow P, Yamada H, Elder JH, Ravkov EV, Nichol ST, Compans RW, Campbell KP, Oldstone MB. Identification of alpha-dystroglycan as a receptor for lymphocytic choriomeningitis virus and Lassa fever virus. Science. 1998;282(5396):2079–81. doi: 10.1126/science.282.5396.2079. PubMed PMID: 9851928.

16. Radoshitzky SR, Abraham J, Spiropoulou CF, Kuhn JH, Nguyen D, Li W, Nagel J, Schmidt PJ, Nunberg JH, Andrews NC, Farzan M, Choe H. Transferrin receptor 1 is a cellular receptor for New World haemorrhagic fever arenaviruses. Nature. 2007;446(7131):92–6. Epub 2007/02/07. doi: 10.1038/nature05539. PubMed PMID: 17287727; PMCID: PMC3197705.

17. Flanagan ML, Oldenburg J, Reignier T, Holt N, Hamilton GA, Martin VK, Cannon PM. New world clade B arenaviruses can use transferrin receptor 1 (TfR1)-dependent and -independent entry pathways, and glycoproteins from human pathogenic strains are associated with the use of TfR1. J Virol. 2008;82(2):938–48. Epub 2007/11/14. doi: 10.1128/JVI.01397-07. PubMed PMID: 18003730; PMCID: PMC2224602.

18. Nunberg JH, York J. The curious case of arenavirus entry, and its inhibition. Viruses. 2012;4(1):83–101. Epub 2012/01/13. doi: 10.3390/v4010083. PubMed PMID: 22355453; PMCID: PMC3280523.

19. Di Simone C, Zandonatti MA, Buchmeier MJ. Acidic pH triggers LCMV membrane fusion activity and conformational change in the glycoprotein spike. Virology. 1994;198(2):455–65. doi: 10.1006/viro.1994.1057. PubMed PMID: 8291229.

20. Wright KE, Salvato MS, Buchmeier MJ. Neutralizing epitopes of lymphocytic choriomeningitis virus are conformational and require both glycosylation and disulfide bonds for expression. Virology. 1989;171(2):417–26. doi: 10.1016/0042-6822(89)90610-7. PubMed PMID: 2474891.

21. Wright KE, Spiro RC, Burns JW, Buchmeier MJ. Post-translational processing of the glycoproteins of lymphocytic choriomeningitis virus. Virology. 1990;177(1):175–83. doi: 10.1016/0042-6822(90)90471-3. PubMed PMID: 2141203; PMCID: PMC7130728.

22. Manning JT, Seregin AV, Yun NE, Koma T, Huang C, Barral J, de la Torre JC, Paessler S. Absence of an N-Linked glycosylation motif in the glycoprotein of the live-attenuated Argentine hemorrhagic fever vaccine, Candid #1, results in Its improper processing, and reduced surface expression. Front Cell Infect Microbiol. 2017;7:20. Epub 2017/02/06. doi: 10.3389/fcimb.2017.00020. PubMed PMID: 28220142; PMCID: PMC5292626.

23. Manning JT, Yun NE, Seregin AV, Koma T, Sattler RA, Ezeomah C, Huang C, de la Torre JC, Paessler S. The glycoprotein of the live-attenuated Junin virus vaccine strain induces endoplasmic reticulum stress and forms aggregates prior to degradation in the lysosome. J Virol. 2020;94(8). Epub 2020/03/31. doi: 10.1128/JVI.01693-19. PubMed PMID: 31996435; PMCID: PMC7108856.

24. Enria DA, Barrera Oro JG. Junin virus vaccines. Curr Top Microbiol Immunol. 2002;263:239–61. doi: 10.1007/978-3-642-56055-2_12. PubMed PMID: 11987817.

25. Barrera Oro JG, McKee KT. Toward a vaccine against Argentine hemorrhagic fever. Bull Pan Am Health Organ. 1991;25(2):118–26. PubMed PMID: 1654168.

26. Albariño CG, Bird BH, Chakrabarti AK, Dodd KA, Flint M, Bergeron E, White DM, Nichol ST. The major determinant of attenuation in mice of the Candid1 vaccine for Argentine hemorrhagic fever is located in the G2 glycoprotein transmembrane domain. J Virol. 2011;85(19):10404–8. Epub 2011/07/27. doi: 10.1128/JVI.00856-11. PubMed PMID: 21795336; PMCID: PMC3196416.

27. Koma T, Patterson M, Huang C, Seregin AV, Maharaj PD, Miller M, Smith JN, Walker AG, Hallam S, Paessler S. Machupo virus expressing GPC of the Candid#1 vaccine strain of Junin virus Is highly attenuated and immunogenic. J Virol. 2016;90(3):1290–7. Epub 2015/11/18. doi: 10.1128/JVI.02615-15. PubMed PMID: 26581982; PMCID: PMC4719595.

28. Koma T, Huang C, Aronson JF, Walker AG, Miller M, Smith JN, Patterson M, Paessler S. The ectodomain of glycoprotein from the Candid#1 vaccine strain of Junin virus rendered Machupo virus partially attenuated in mice lacking IFN-αβ/γ receptor. PLoS Negl Trop Dis. 2016;10(8):e0004969. Epub 2016/08/31. doi: 10.1371/journal.pntd.0004969. PubMed PMID: 27580122; PMCID: PMC5006991.

29. Koma T, Huang C, Coscia A, Hallam S, Manning JT, Maruyama J, Walker AG, Miller M, Smith JN, Patterson M, Abraham J, Paessler S. Glycoprotein N-linked glycans play a critical role in arenavirus pathogenicity. PLoS Pathog. 2021;17(3):e1009356. Epub 20210301. doi: 10.1371/journal.ppat.1009356. PubMed PMID: 33647064; PMCID: PMC7951981.

30. Patterson M, Koma T, Seregin A, Huang C, Miller M, Smith J, Yun N, Paessler S. A substitution in the transmembrane region of the glycoprotein leads to an unstable attenuation of Machupo virus. J Virol. 2014;88(18):10995–9. Epub 2014/07/16. doi: 10.1128/JVI.01007-14. PubMed PMID: 25031335; PMCID: PMC4178861.

31. Patterson M, Seregin A, Huang C, Kolokoltsova O, Smith J, Miller M, Yun N, Poussard A, Grant A, Tigabu B, Walker A, Paessler S. Rescue of a recombinant Machupo virus from cloned cDNAs and in vivo characterization in interferon (αβ/γ) receptor double knockout mice. J Virol. 2014;88(4):1914–23. Epub 2013/11/27. doi: 10.1128/JVI.02925-13. PubMed PMID: 24284323; PMCID: PMC3911560.

32. Sommerstein R, Flatz L, Remy MM, Malinge P, Magistrelli G, Fischer N, Sahin M, Bergthaler A, Igonet S, Ter Meulen J, Rigo D, Meda P, Rabah N, Coutard B, Bowden TA, Lambert PH, Siegrist CA, Pinschewer DD. Arenavirus glycan shield promotes neutralizing antibody evasion and protracted infection. PLoS Pathog. 2015;11(11):e1005276. Epub 2015/11/20. doi: 10.1371/journal.ppat.1005276. PubMed PMID: 26587982; PMCID: PMC4654586.

33. Watanabe Y, Raghwani J, Allen JD, Seabright GE, Li S, Moser F, Huiskonen JT, Strecker T, Bowden TA, Crispin M. Structure of the Lassa virus glycan shield provides a model for immunological resistance. Proc Natl Acad Sci U S A. 2018;115(28):7320–5. Epub 2018/06/25. doi: 10.1073/pnas.1803990115. PubMed PMID: 29941589; PMCID: PMC6048489.

34. Golden JW, Beitzel B, Ladner JT, Mucker EM, Kwilas SA, Palacios G, Hooper JW. An attenuated Machupo virus with a disrupted L-segment intergenic region protects guinea pigs against lethal Guanarito virus infection. Sci Rep. 2017;7(1):4679. Epub 2017/07/05. doi: 10.1038/s41598-017-04889-x. PubMed PMID: 28680057; PMCID: PMC5498534.

35. Bell TM, Bunton TE, Shaia CI, Raymond JW, Honnold SP, Donnelly GC, Shamblin JD, Wilkinson ER, Cashman KA. Pathogenesis of Bolivian hemorrhagic fever in guinea pigs. Vet Pathol. 2016;53(1):190–9. Epub 2015/07/02. doi: 10.1177/0300985815588609. PubMed PMID: 26139838.

36. Mantlo E, Paessler S, Huang C. Differential immune responses to hemorrhagic fever-causing arenaviruses. Vaccines (Basel). 2019;7(4). Epub 2019/10/05. doi: 10.3390/vaccines7040138. PubMed PMID: 31581720; PMCID: PMC6963578.

37. Weissenbacher MC, Avila MM, Calello MA, Merani MS, McCormick JB, Rodriguez M. Effect of ribavirin and immune serum on Junin virus-infected primates. Med Microbiol Immunol. 1986;175(2-3):183–6. PubMed PMID: 3014292.

38. Brouillette RB, Phillips EK, Ayithan N, Maury W. Differences in Glycoprotein Complex Receptor Binding Site Accessibility Prompt Poor Cross-Reactivity of Neutralizing Antibodies between Closely Related Arenaviruses. J Virol. 2017;91(7). Epub 20170313. doi: 10.1128/JVI.01454-16. PubMed PMID: 28100617; PMCID: PMC5355595.

39. Zeltina A, Krumm SA, Sahin M, Struwe WB, Harlos K, Nunberg JH, Crispin M, Pinschewer DD, Doores KJ, Bowden TA. Convergent immunological solutions to Argentine hemorrhagic fever virus neutralization. Proc Natl Acad Sci U S A. 2017;114(27):7031–6. Epub 20170619. doi: 10.1073/pnas.1702127114. PubMed PMID: 28630325; PMCID: PMC5502616.

40. Bonhomme CJ, Capul AA, Lauron EJ, Bederka LH, Knopp KA, Buchmeier MJ. Glycosylation modulates arenavirus glycoprotein expression and function. Virology. 2011;409(2):223–33. Epub 2010/11/05. doi: 10.1016/j.virol.2010.10.011. PubMed PMID: 21056893; PMCID: PMC3053032.

41. Saphire EO, Schendel SL, Fusco ML, Gangavarapu K, Gunn BM, Wec AZ, Halfmann PJ, Brannan JM, Herbert AS, Qiu X, Wagh K, He S, Giorgi EE, Theiler J, Pommert KBJ, Krause TB, Turner HL, Murin CD, Pallesen J, Davidson E, Ahmed R, Aman MJ, Bukreyev A, Burton DR, Crowe JE, Davis CW, Georgiou G, Krammer F, Kyratsous CA, Lai JR, Nykiforuk C, Pauly MH, Rijal P, Takada A, Townsend AR, Volchkov V, Walker LM, Wang CI, Zeitlin L, Doranz BJ, Ward AB, Korber B, Kobinger GP, Andersen KG, Kawaoka Y, Alter G, Chandran K, Dye JM, Consortium VHFI. Systematic analysis of monoclonal antibodies against Ebola virus GP defines features that contribute to protection. Cell. 2018;174(4):938–52.e13. doi: 10.1016/j.cell.2018.07.033. PubMed PMID: 30096313; PMCID: PMC6102396.

42. Ko YA, Yu YH, Wu YF, Tseng YC, Chen CL, Goh KS, Liao HY, Chen TH, Cheng TR, Yang AS, Wong CH, Ma C, Lin KI. A non-neutralizing antibody broadly protects against influenza virus infection by engaging effector cells. PLoS Pathog. 2021;17(8):e1009724. Epub 20210805. doi: 10.1371/journal.ppat.1009724. PubMed PMID: 34352041; PMCID: PMC8341508.

43. Henry Dunand CJ, Leon PE, Huang M, Choi A, Chromikova V, Ho IY, Tan GS, Cruz J, Hirsh A, Zheng NY, Mullarkey CE, Ennis FA, Terajima M, Treanor JJ, Topham DJ, Subbarao K, Palese P, Krammer F, Wilson PC. Both Neutralizing and Non-Neutralizing Human H7N9 Influenza Vaccine-Induced Monoclonal Antibodies Confer Protection. Cell Host Microbe. 2016;19(6):800–13. doi: 10.1016/j.chom.2016.05.014. PubMed PMID: 27281570; PMCID: PMC4901526.

44. Hangartner L, Zellweger RM, Giobbi M, Weber J, Eschli B, McCoy KD, Harris N, Recher M, Zinkernagel RM, Hengartner H. Nonneutralizing antibodies binding to the surface glycoprotein of lymphocytic choriomeningitis virus reduce early virus spread. J Exp Med. 2006;203(8):2033–42. Epub 2006/07/31. doi: 10.1084/jem.20051557. PubMed PMID: 16880253; PMCID: PMC2118372.

45. Straub T, Schweier O, Bruns M, Nimmerjahn F, Waisman A, Pircher H. Nucleoprotein-specific nonneutralizing antibodies speed up LCMV elimination independently of complement and FcγR. Eur J Immunol. 2013;43(9):2338–48. Epub 20130715. doi: 10.1002/eji.201343565. PubMed PMID: 23749409.

46. Rodriguez-Carreno MP, Nelson MS, Botten J, Smith-Nixon K, Buchmeier MJ, Whitton JL. Evaluating the immunogenicity and protective efficacy of a DNA vaccine encoding Lassa virus nucleoprotein. Virology. 2005;335(1):87–98. doi: 10.1016/j.virol.2005.01.019. PubMed PMID: 15823608.

47. Abreu-Mota T, Hagen KR, Cooper K, Jahrling PB, Tan G, Wirblich C, Johnson RF, Schnell MJ. Non-neutralizing antibodies elicited by recombinant Lassa-Rabies vaccine are critical for protection against Lassa fever. Nat Commun. 2018;9(1):4223. Epub 20181011. doi: 10.1038/s41467-018-06741-w. PubMed PMID: 30310067; PMCID: PMC6181965.

48. Kenyon RH, Condie RM, Jahrling PB, Peters CJ. Protection of guinea pigs against experimental Argentine hemorrhagic fever by purified human IgG: importance of elimination of infected cells. Microb Pathog. 1990;9(4):219–26. doi: 10.1016/0882-4010(90)90010-n. PubMed PMID: 1965845.

49. Droniou-Bonzom ME, Reignier T, Oldenburg JE, Cox AU, Exline CM, Rathbun JY, Cannon PM. Substitutions in the glycoprotein (GP) of the Candid#1 vaccine strain of Junin virus increase dependence on human transferrin receptor 1 for entry and destabilize the metastable conformation of GP. J Virol. 2011;85(24):13457–62. Epub 2011/10/07. doi: 10.1128/jvi.05616-11. PubMed PMID: 21976641; PMCID: PMC3233171.

50. York J, Nunberg JH. Epistastic interactions within the Junín virus envelope glycoprotein complex provide an evolutionary barrier to reversion in the live-attenuated Candid#1 vaccine. J Virol. 2018;92(1). Epub 2017/10/27. doi: 10.1128/jvi.01682-17. PubMed PMID: 29070682; PMCID: PMC5730776.

51. York J, Nunberg JH. Intersubunit interactions modulate pH-induced activation of membrane fusion by the Junin virus envelope glycoprotein GPC. J Virol. 2009;83(9):4121–6. Epub 20090218. doi: 10.1128/JVI.02410-08. PubMed PMID: 19224989; PMCID: PMC2668491.

52. Gowen BB, Hickerson BT, York J, Westover JB, Sefing EJ, Bailey KW, Wandersee L, Nunberg JH. Second-generation live-attenuated Candid#1 vaccine virus resists reversion and protects against lethal Junín virus infection in guinea pigs. J Virol. 2021. Epub 2021/05/05. doi: 10.1128/JVI.00397-21. PubMed PMID: 33952638.

53. Bonhomme CJ, Knopp KA, Bederka LH, Angelini MM, Buchmeier MJ. LCMV glycosylation modulates viral fitness and cell tropism. PLoS One. 2013;8(1):e53273. Epub 2013/01/07. doi: 10.1371/journal.pone.0053273. PubMed PMID: 23308183; PMCID: PMC3538765.

54. Emonet SF, Seregin AV, Yun NE, Poussard AL, Walker AG, de la Torre JC, Paessler S. Rescue from cloned cDNAs and in vivo characterization of recombinant pathogenic Romero and live-attenuated Candid #1 strains of Junin virus, the causative agent of Argentine hemorrhagic fever disease. J Virol. 2011;85(4):1473–83. Epub 2010/12/01. doi: 10.1128/JVI.02102-10. PubMed PMID: 21123388; PMCID: PMC3028888.

